# Small molecules targeting the structural dynamics of AR-V7 partially disordered protein using deep learning and physics based models

**DOI:** 10.1101/2024.02.23.581804

**Authors:** Pantelis Karatzas, Z. Faidon Brotzakis, Haralambos Sarimveis

## Abstract

Partially disordered proteins can contain both stable and unstable secondary structure segments and are involved in various (mis)functions in the cell. The extensive conformational dynamics of partially disordered proteins scaling with extent of disorder and length of the protein hampers the efficiency of traditional experimental and in-silico structure-based drug discovery approaches. Therefore new efficient paradigms in drug discovery taking into account conformational ensembles of proteins need to emerge. In this study, using as a test case the AR-V7 transcription factor splicing variant related to prostate cancer, we present an automated methodology that can accelerate the screening of small molecule binders targeting partially disordered proteins. By swiftly identifying the conformational ensemble of AR-V7, and reducing the dimension of binding-sites by a factor of 90 by applying appropriate physicochemical filters, we combine physics based molecular docking and multi-objective classification machine learning models that speed up the screening of thousands of compounds targeting AR-V7 multiple binding sites. Our method not only identifies previously known binding sites of AR-V7, but also discovers new ones, as well as increases the multi-binding site hit-rate of small molecules by a factor of 10 compared to naive physics-based molecular docking.

## 1 Introduction

The process of discovering new drugs is both expensive and time-consuming, facing several hurdles including the challenge of finding at the preclinical level effective drug candidates through high-throughput screening methods, which often have low success rates^1,2^. To tackle these issues, the use of computer-aided drug discovery (CADD) has become increasingly important. CADD can make the drug discovery process faster and more successful^3,4^. A key technique in CADD is molecular docking, an energy based technique which allows to quickly assess the affinity of millions of compounds such as small molecules and peptides against various drug targets, whose three-dimensional shapes is a-priori known.

However, the application of molecular docking in drug discovery encounters unique challenges when targeting partially disordered proteins (PDPs). PDPs, characterized by a mix of stable secondary-tertiary structured and disordered segments with only transient secondary structure formed^1,2,5^. Due to this interplay, PDPs can be understood in terms of a structural ensemble, that is the collection of structures equipped with the underlying populations according to thermodynamics, and cannot be characterized by a single conformation alone, thereby rendering conventional drug discovery techniques not suitable for the design of a small molecule. Such conformations often lack a stable and single active binding site that can be used for in-silico screening and designing small molecules.

Despite these obstacles, targeting PDPs is a critical area of research due to their significant role in various biological processes^6^ and their link to a multitude of human diseases, such as cancer^2^, neurodegenerative diseases^3^ and many others^6^. Previous analyses of the structures of 51 intrinsically disordered proteins (IDPs), which are PDPs that have no stable secondary structure fold at all, revealed that on average, PDPs contain 50% more druggable pockets than fully structured proteins. Moreover, the average probability of finding druggable sites was approximately 9 %, compared to just 5 % in structured proteins^7^. Since partially disordered proteins encompass both structured and disordered regions, and considering that they form a broader category than IDPs, it is reasonable to infer that PDPs might similarly exhibit a high potential for druggability. However, PDPs have not been as extensively studied in this context, suggesting the need for further research to understand their full druggability potential. Currently, there is no experimental technique available that can directly trace the atomistic structural ensemble of partially disordered proteins^8^. Integrative structural biology methods requiring SAXS or NMR data as input into physics based molecular dynamics have greatly helped to determine structural ensembles of partially disordered proteins^9,10^. AlphaFold has revolutionized protein structure prediction^11^. More evidence supports that information about conformational dynamics is contained in AlphaFold^12–14^. Current advances in structural ensemble prediction recently showed how to introduce AlphaFold data into physics based molecular dynamics in order to infer structural ensembles of partially disordered proteins in a time-efficient manner^12^. However, the challenge persists in the molecular screening process for both determining binding sites and screening of molecules, since a binding site can have numerous conformations. Throughout the various conformational states of a PDP, multiple different residues can form multiple structurally different pockets, thereby complicating the selection of a single binding site. In this study, we present a novel methodology designed to refine the identification of binding sites on PDPs and to accelerate the screening of molecular libraries using deep learning models. This approach leverages multiple criteria applied to the conformations of partially disordered proteins to identify sites with favorable characteristics for drug interaction. The computational workflow in this study begins with the retrieval of the conformational ensemble of the target biomolecule, by utilizing AlphaFold-MetaInference due to its efficiency in speed and accuracy. Following this, we focus on the detection and selection of binding sites, which varies according to the protein’s biological function. The next phase involves molecular docking, using a set of molecules randomly sampled from the ChEMBL database, although other criteria like Lipinski’s rule of five could be employed for more effective sub-sampling of the chemical space. The data generated from this docking stage is then used to create a multi-label classification Quantitative Structure-Activity Relationship (QSAR) model, which accelerates the screening process by allowing broader inference of the chemical space. In particular, molecules predicted as active in most of the docking sites and selected for further analysis and docking simulations.

The proposed methodology is applied to a pertinent case study, that of nuclear hormone receptors, such as the androgen receptor (AR) transcription factor, a category of proteins considered largely undraggable^11^. AR, is characterized by a structured ligand-binding domain (LBD), and therapies targeting this domain are commonly used as the first line of treatment for AR-driven prostate cancer^15,16^. However, approximately 20 % of prostate cancer patients advance to a more lethal stage known as castration-resistant prostate cancer (CRPC). This progression is often marked by the emergence of constitutively active AR splicing variants such as AR-V7. These CRPC-associated AR variant is devoid of the LBD and consists solely of the DNA-binding domain and an intrinsically disordered activation domain (AD), making it resistant to treatments targeting the LBD^8,17^. In particular AR-V7 splicing variant undergoes phase separation caused by interactions between sticky residues thereby promoting its transcriptional activity leading to tumor growth^18^. The hypothesis of this study is that by identifying efficiently small molecule binders at multiple binding pockets around sticky residues in the structural ensemble of AR-V7 can potentially inhibit phase separation leading to tumor growth. Overall, the structural-ensemble based small molecule binder optimization against these disease relevant partially disordered activation domains in AR-V7 provides a crucial insight into the challenges of targeting PDPs in drug discovery.

## 2 Results

In this study, we focus on unraveling the complexities of partially disordered proteins (PDPs) and their significant role in drug discovery. We introduce a novel methodology tailored to enhance the efficiency of screening molecular libraries, with particular emphasis on their interaction with PDPs. Central to our approach is a comprehensive analysis of PDP structural ensembles, which is essential for understanding the dynamic and diverse nature of their potential binding sites. A key aspect of our methodology is the use of molecular docking data to train a multi-label classification model. This model is designed to rapidly predict the interaction potential of compounds across all relevant PDP binding sites, each with its distinct conformational state. By doing so, we are able to screen large libraries of molecules efficiently against multiple relevant binding sites, significantly accelerating the identification of promising compounds. This process not only aids in a deeper understanding of PDPs’ druggability but also facilitates the discovery of novel therapeutic agents for diseases where PDPs play a pivotal role.

The proposed methodology consists of the following steps:

1. Retrieve the conformational ensemble of the biomolecular target of interest. For this purpose, AlphaFold-Metainference ^12^ is employed, due to its speed and accuracy.
2. Pocket detection and selection of binding sites. This part of the methodology is tailored to each protein, depending on the biological function of the protein that needs to be altered. For instance, in targeting AR-V7, to prevent abnormal liquid-liquid phase separation, - a phenomenon that leads to aberrant transcription activation and tumor growth-our focus is on detecting binding sites around ‘sticky’ residues, as these are responsible for initiating liquid-liquid phase separation.
3. Selection of a list of small molecules for molecular docking against the detected cavities. In our case we sampled randomly molecules from the Chembl database. However, other criteria, such as the Lipinski rule of 5, could also be applied for more effective subsampling of the chemical space.
4. Use the data created in step 3, to develop a multi-label classification QSAR model for predicting the activity against all binding sites selected in step 2. This model is designed to accelerate the screening process by enabling the inference of a broader chemical space. In our specific scenario, molecules that are predicted as ‘active’ (labeled as ‘1’) in over 70% of the docking sites are considered promising candidates. It’s important to note that this threshold can vary depending on the protein being studied and the results of the screening.
5. Re-evaluate with docking molecules meeting this criterion.

### 2.1 AR-V7 structural ensemble

The structural ensemble of the prostate cancer relevant AR-V7 splicing variant, generated by AF-MI is revealed in Fig. 1A. AR-V7 exhibits remarkable heterogeneity of structures, especially in corresponding to the AD region of AR-V7, which well captures current understanding of the IDR part of the AD domain^18,19,19^. More explicit secondary structure analysis shown in Fig. 1B reveals high propensity secondary structure regions (helicity ¿60 %) around residues 56-80 (polyQ domain), 237-244, 578-588 and 614-623 (614SCRLRK-CYEAG623). Transient helices (2%¡helicity¡10%) form at residues 23-27 (the 23FQNLF27 segment),173-198, 397-402 (the 397SAWAAA402 segment) and 634-636(634GNC636). The positions of the helical residues majorly agree with NMR and MD findings by Basu et. al. ^20^. High beta-sheet propensity shown in Fig. 1C is exhibited only at residues 566-577 of the DBD domain and high propensity of coil regions is manifested in the rest of the sequence as shown in Fig. 1D. Since AR-V7 undergoes phase separation which in term enhances transcriptional activity^20^, we embarked to quantify the sticky residues using as proxy the residue based root mean square fluctuation quantified by using the structural ensemble. Residues of regions tau1, tau5* and tau5 exhibits a distinct lower flexibility (rmsf¡4.5 nm2), pinpointing hindrance and inter-intra molecular interactions of these regions. Evidently, the tau-5 region contains more sticky residues, in agreement with 1H-15 N NMR^20^.

**Figure 1:**
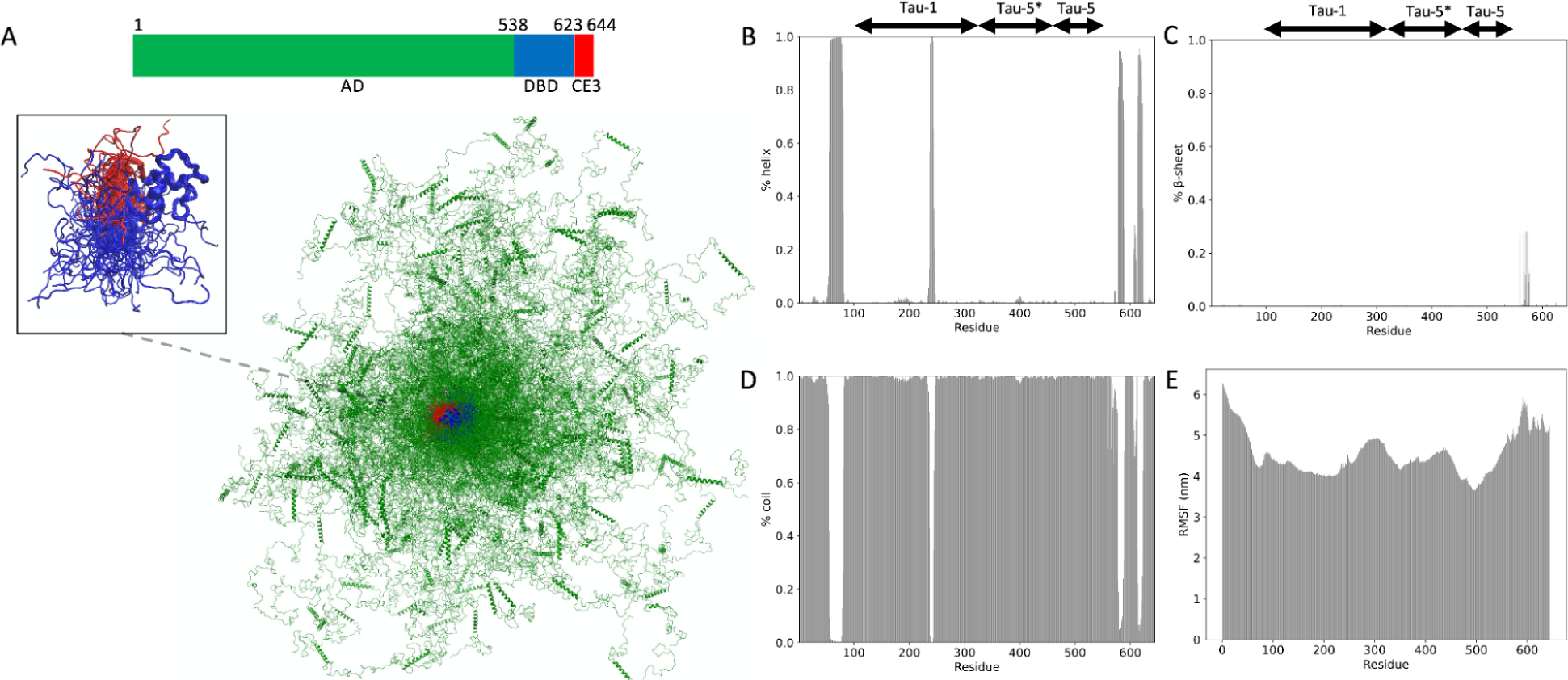
Structural ensemble A) Primary sequence and structural ensemble of AR-V7. AD, DBD and CE3 domains are colored in green, blue and red respectively. B,C,D) Residue based population fraction of *α*-helix, *β*-sheet and coil secondary structure. E) Residue based Root Mean Square Fluctuation. We applied our methodology to AR-V7, a variant of the androgen receptor associated with prostate cancer. This application led to the identification of several compounds that demonstrated high affinity for specific binding sites. These findings, stemming from our efficient screening process and the predictions of our multi-label QSAR model, underscore the efficacy of our approach in identifying potential therapeutic agents for complex PDPs such as AR-V7. The detailed results are presented in the following sections:

### 2.2 Binding site selection

Filtering and selecting the appropriate binding sites is critical in our proposed pipeline. Before filtering, 3718 cavities were detected across the ensemble Fig. 2A), with each conformation in the ensemble displaying multiple binding sites. Molecular docking against all of them is computationally prohibitive. As noted in the methods section, by focusing on sticky residues and large cavities as quantified by Root Mean Squared Fluctuation (RMSF), hydropathy and volume/area of cavities, we narrowed down our selection to 41 binding sites (see Fig. 2B). This reduction in binding sites was instrumental in decreasing the computational cost of docking calculations by 90-fold. Further analysis of the residue identity of these 41 binding sites is presented in Fig. 2C where we categorize them into 26 distinct binding site groups based on common residue sharing more than 60%. Binding site clusters 0-6 contain multiple members, while the remaining 19 clusters consist of single members (see Fig. 2D). Fig. 2C shows that the most of the binding sites are located in regions tau-5 (19 binding sites), followed by tau-1 (17 binding sites) and tau-5*/tau-5 (5 binding sites). This distribution is significant, particularly because regions tau-1 (around residue 180) and tau-5* (397-402) are known for transient helix formation and have been shown to inhibit phase separation and transcriptional activity of AR-V7 when targeted by small molecule binders, especially around residues 393, 397, 437, and 445^20^. Our binding sites with IDs 2,9,18 contain such critical residues. In Fig. 2E,F we show two example configurations where the binding sites are located in the tau-1 and tau-5*and tau-5 region of the AD domain, corresponding to binding site IDs 0 (spanning tau-5 region),1(tau-5 region), 2 (spanning regions tau-5*,tau-5).

**Figure 2:**
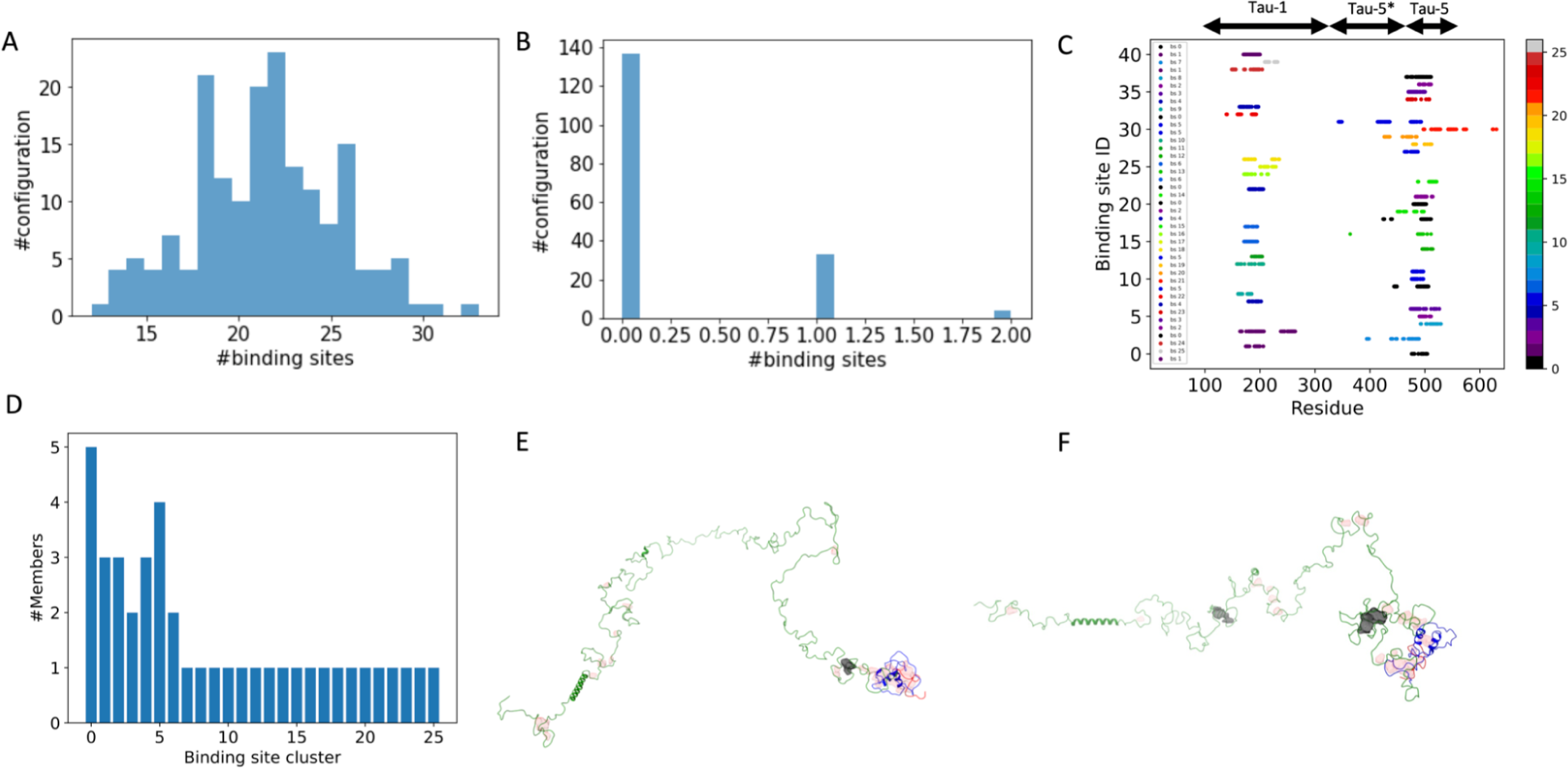
Binding site selection. A) Distribution of all binding sites throughout the ensemble B) Distribution of filtered binding sites throughout the ensemble. C) Residue based identity of binding sites. Colors correspond to each binding cluster. D) Cluster analysis of binding sites. E,F) Two example conformations. with red are illustrated all binding sites. With black the ones that pass the binding site selection filters. The filtered binding sites comprise residues E) 476-477, 479-81, 488 493-498,501-504 (tau-5 region), F) 174-179, 181, 190-200, 206 (tau-1 region) and 394-396. 398, 439-441,447-449, 460-462, 470-479, 482-489 (tau-5*/tau-5 region).

**Figure 3:**
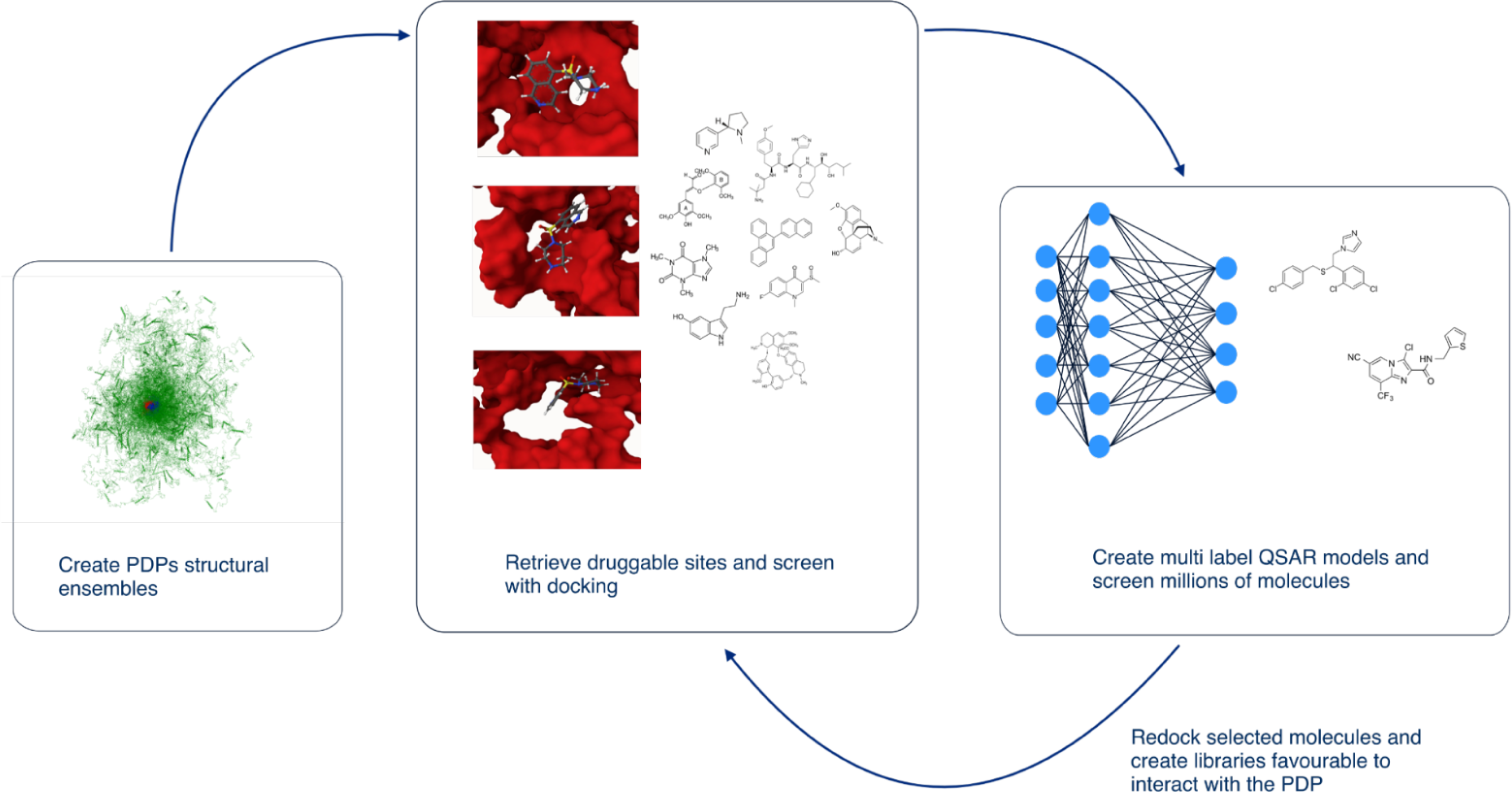
Structural ensemble based drug discovery using physics and AI models. Multilabel Retrieve druggable docking sites from structural ensembles, dock molecules, create multi-label QSAR models and screen billions of molecules. Selected molecules are re docked and prioritized.

### 2.3 Molecular Docking

Deep Dock and the Deep Docking protocol^21^ have been established to expedite the screening of molecules against a single docking site, yielding excellent results in terms of both time reduction and the enrichment of potential molecular hits. However, as previously discussed, the nature of partially disordered proteins (PDPs) differs from the structured regions of the proteome, often presenting multiple docking sites when targeting a PDP. This presents a challenge in the scalability of screening vast numbers of molecules; for example, in the case of the AR-V7 protein with its 41 selected docking sites, screening 1 million molecules would result in a total of 41 million screenings – a task not computationally efficient. Consequently, the careful selection of an initial pool of molecules was crucial to achieve the best possible outcomes. For this purpose, a list of 5640 compounds was randomly selected from the Chembl database^22^. The binding energies of these 5640 compounds against all 41 docking sites were computed using Autodock Vina, with more details on this process provided in the Methods section.

### 2.4 Development of a Deep Learning multi-label classification model for predicting binding potential to all AR-V7 41 binding sites

To efficiently screen a wide array of compounds, we utilized the molecular docking data to train a multi-label classification QSAR model. This model predicts a compound’s activity against any of 41 binding sites. The model’s dataset was created by first establishing a binding energy threshold for each site, defined as the highest energy level within the top 5% of compounds with the lowest binding energies. Compounds were then classified as active (”1”) if their docking energy was below this threshold, and inactive (”0”) otherwise, resulting in each compound having 41 potential classification outcomes. To counter the heavy imbalance toward inactive molecules in the dataset, we focused on training the model with the top 5% of binding energies by oversampling active molecules. The model thus trained was then utilized to screen molecular libraries. Details about the QSAR modeling process are provided in the “Methods” section.

Machine Learning and structural dynamics based optimization of small molecules against AR-V7. The screening process for the AR-V7 protein, with its 41 docking sites, demands significant computational power and resources. When conducting screenings against such a number of docking sites, or in the case of blind docking, the calculations required an increase in direct proportion to the number of protein binding sites — in this instance, by a factor of 41. Although various methods and protocols^21,23^ incorporate machine learning models to streamline this process, only a few computational methodologies are tailored specifically for PDPs or their regions. Notably, our proposed method enabled us to rapidly screen a library of almost 2 million molecules, randomly selected from the ChEMBL database. Out of this extensive screening, we identified molecules that were active against at least 70% of the binding sites, narrowing it down to just 402 compounds, which represents only 0.021% of the total screened molecules. The binding energies of these 402 compounds against all 41 docking sites were calculated using Autodock Vina. Comprehensive results are provided in the supplementary information file screened results.csv, which includes the SMILES representations of these 402 molecules, as well as their calculated binding energies for the 41 docking sites.

To further elucidate the effectiveness of our proposed screening strategy, we carried out a comparative analysis between the 5640 molecules, which were randomly selected for training the multi-label classification model (referred to as ‘naive docking’), and the 402 compounds identified through our screening process (referred to as ‘screened docking’). The results are summarized in Fig. 5 which displays the number of molecules predicted to be active against one to 41 binding sites for both sets of molecules. Notably, only 57/5639 (1%) of the molecules in the naive docking set were predicted to be active against more than 30 binding sites, a figure that increased to over 70/401 (17.5%) in the screened docking set.

**Figure 4:**
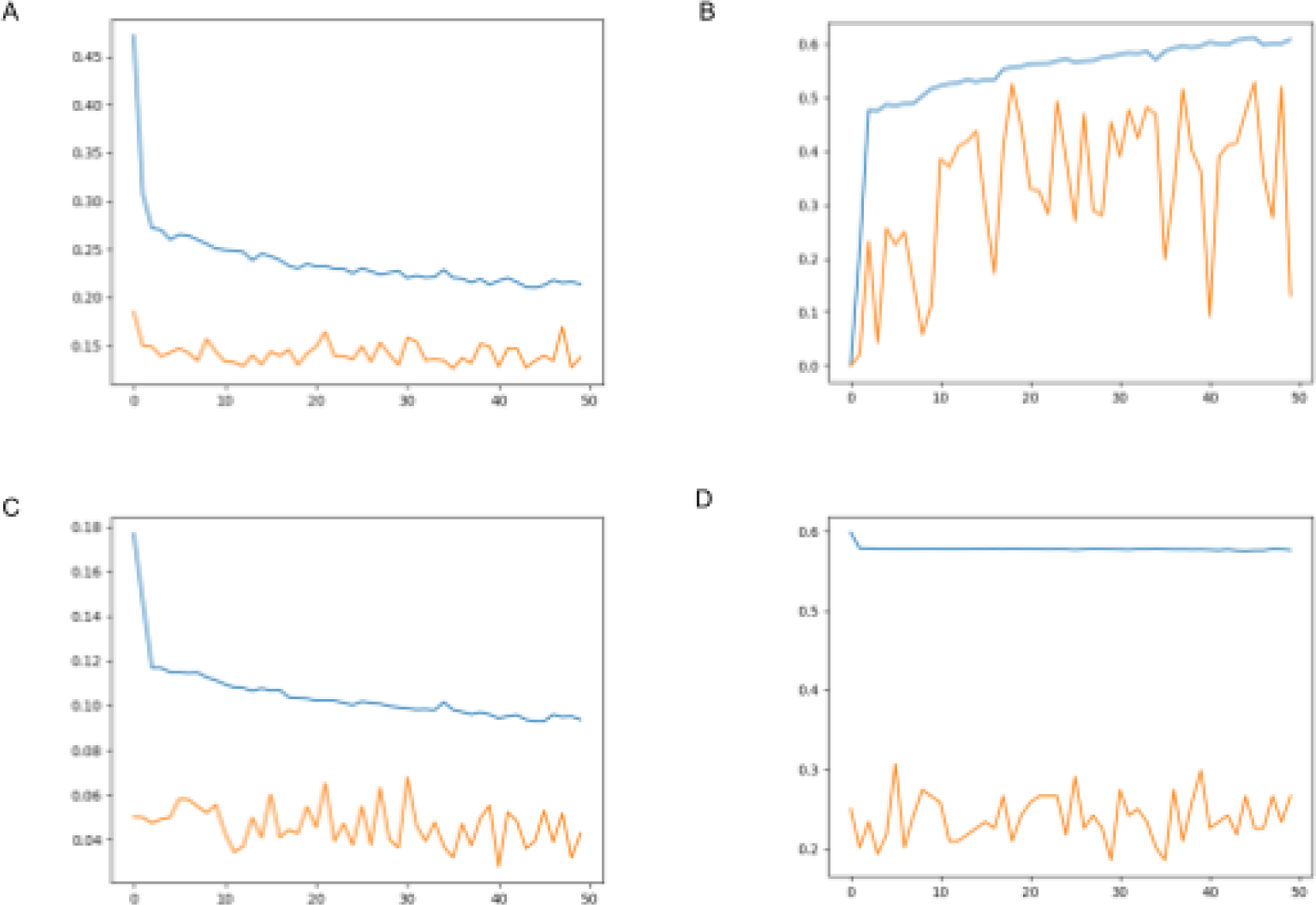
Training the model. A) Binary cross entropy loss per epoch for train and test, B) Average MCC per epoch, C) Hamming loss per epoch, D) Zero one loss per epoch for the top 5% of the docking energies. Blue line corresponds to train set and red to test set.

**Figure 5:**
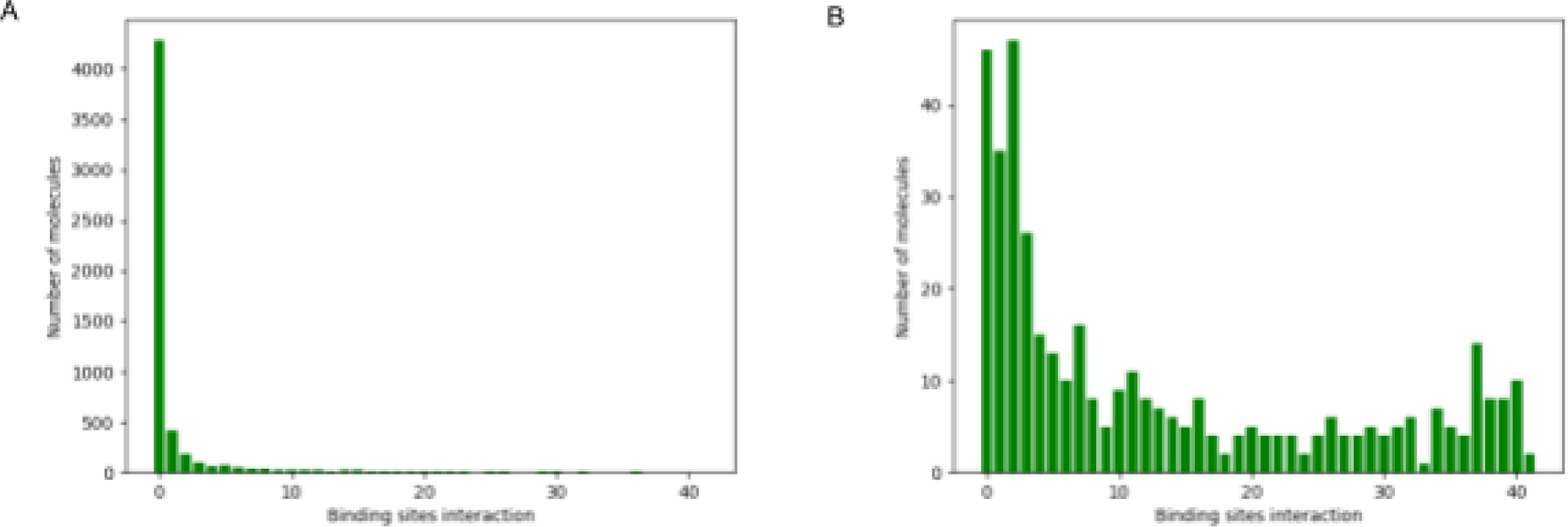
Multibinding site hit rate increase A) Number of “active” molecules with plain docking and top 5%, B) Number of actives on docking site screened against the original threshold.

In addition, we calculated the average binding energies for both sets of molecules and visualized these distributions by plotting their kernel density functions (Fig. 6). The molecules identified through our screening process showed significantly lower average binding energies compared to those in the naive docking set. These results indicate a greater potential for identifying effective hits for targeting the AR-V7 protein among the screened compounds, thereby highlighting the efficiency of our approach in discovering hits against PDPs like AR-V7.

**Figure 6:**
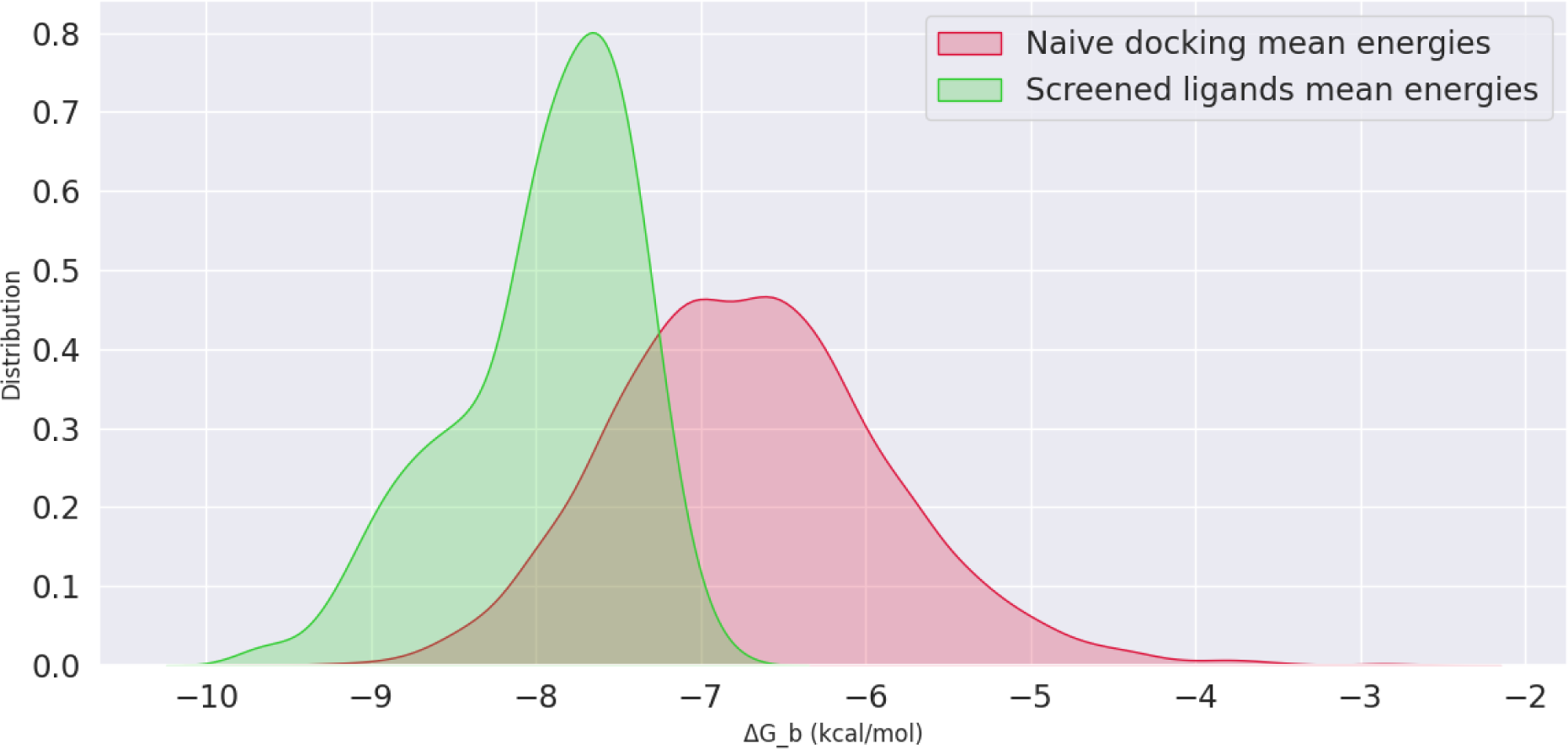
Distribution of binding energies. Kernel density estimation plot for the energies of naive docking and screened molecules

## 3 Methods

### 3.1 Structural ensemble of AR-V7

In order to generate the structural ensemble we carried on the protocol described in Ref.^12^. In particular, we start from the sequence of AR-V7 splicing variant, relevant to prostate cancer^24^. AR-V7 mRNA retains the first three canonical exons, followed by variant-specific cryptic exon 3 (CE3). A splicing event at CE3 leads to an LBD-truncated AR-V7. Hence the sequence of the prostate relevant AR-V7 splicing variant, henceforth mentioned as AR-V7 corresponds to the activation domain (AD), the DNA binding domain (DBD) and a CE3 domain. By using the AR-V7 sequence, listed in table S1 and AlphaFold, we predict the structure of AR-V7 and the corresponding AlphaFold predicted distances di,j. AlphaFold-Metainference (AF-MI)^12^ is used to generate the structural ensemble of AR-V7, by combining the physics based coarse-grained CALVADOS-2^25^ molecular dynamics (MD) model with the AlphaFold predicted distances which are used as restraints in the Metainference framework^26^. However, as in Ref.^25^, we do not consider all di, j distances as restraints but rather a subset. First, since, CALVADOS-2 is an IDP trained coarse-grained model, optimized to reproduce structures of disordered regions, rather than ordered ones, we use the predicted local distance difference test (pLDDT) score of AlphaFold to define regions with pLDDT score ¿0.75 as structured regions. For AR-V7 these regions comprise residues 53-81, 235-244 and 557-628. Such regions are restrained to the AlphaFold-predicted structure by using a root mean square deviation (RMSD) potential and the intra-residue distances corresponding to these structured regions are excluded from the distance restraints. For all other inter-residue distances, we consider only distances that have a Predicted Alignment Error (PAE) lower than 4A (Fig. 7A). For the MD setup part of AF-MI, we set a 5 fs time-step, temperature at 298 K and perform AF-MI simulation in the NVT ensemble for 106 steps per replica with a total of six replicas and frames are saved every 15 ps. To accelerate the sampling we use a Parallel bias metadynamics ^27^ potential along 4 collective variables (Rg1,Rg2,Rg3,Rg4) representing the radius of gyration of the disordered regions 1-52,82-234,245-556,629-644, with original hill. Equivalently to the protocol in^12^, we then used the PULCHRA software^28^ to backmap to atomistic representations of the structures in the structural ensemble. The input files for AF-Mi as well as the AR-V7 ensembles can be found in [our Github or Zenobo]. We have subsampled the ensemble by weights in order to generate an unbiased atomistic AR-V7 ensemble of 350 structures.

**Figure 7:**
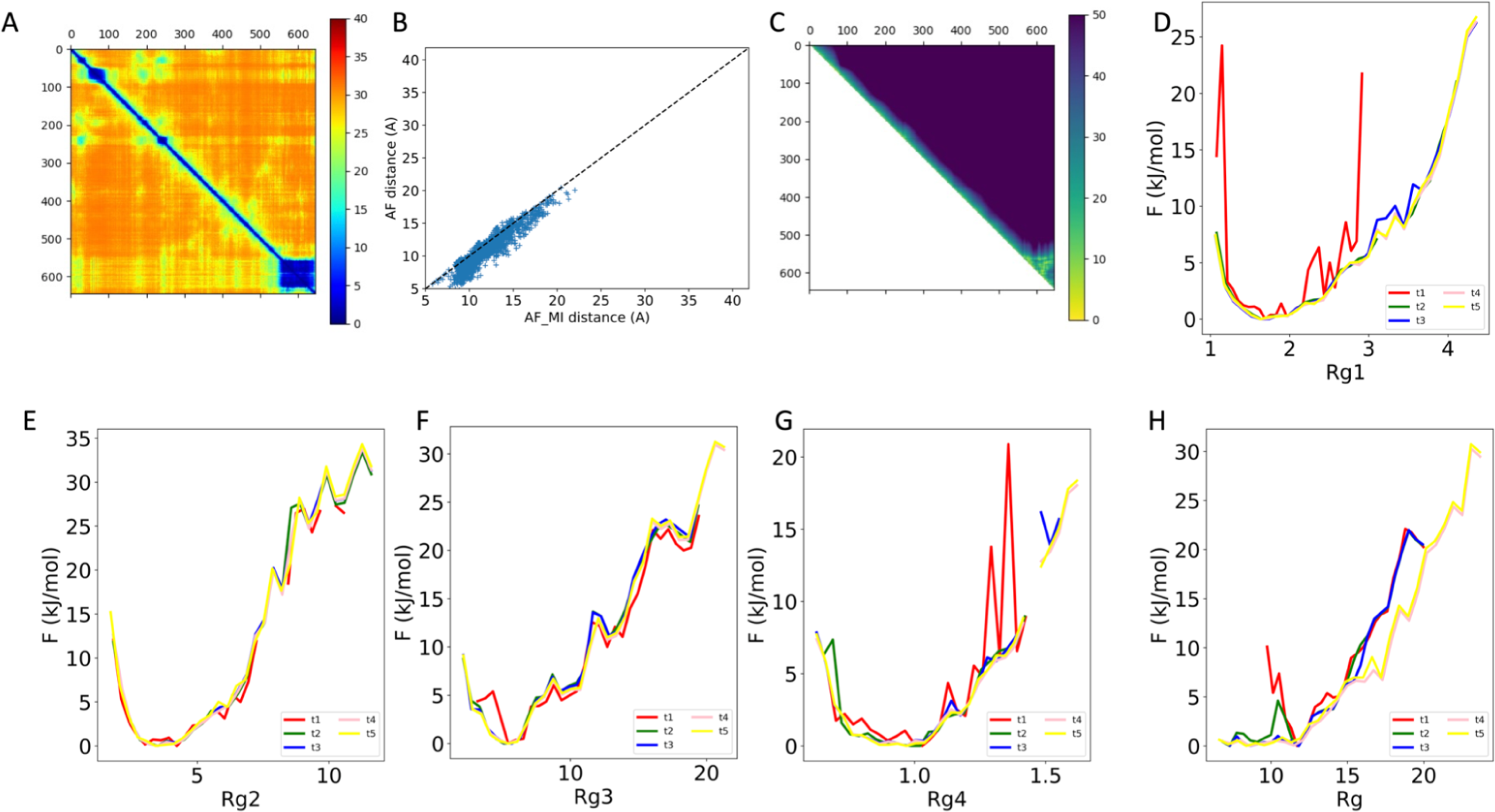
AlphaFold Metainference statistics. A) Residue based predicted alignment error in Angstong. B) AlphaFold predicted and AF-MI restrained distances. C) residue pairwise distance map of the AR-V7 structural ensemble. D,E,F,G) Time dependent free energies of radii of gyration of the disordered segments of AR-V7 (Rg1,Rg2,Rg3,Rg4). H) Time dependent free energy of the radius of gyration of the entire AR-V7 protein.

### 3.2 Pocket detection

To detect and analyze binding sites on the structural ensemble of AR-V7, we use pyKVFinder^29^, an integrable python package that can swiftly detect cavities using as input, protein pdb files and characterize them according to different filters such as hydropathy, volume, number of aliphatic apolar, aromatic, polar uncharged, negatively charged and positively charged amino acids in a cavity. Such physicochemical characterisation of the cavities informs the selection of the favorable binding sites for the purpose of in-silico molecular docking. For the purpose of binding site selection, we require that it contains more than 5 residues, and abides to an upper hydropathy filter of 0, a lower/upper area of 80/2480 A2 and a lower volume of 120 A3only the hydrophobic ones and relatively big to be able to screen relatively big optimizable molecules. On top of these filters, we add a third filter, namely that the RMSF of the residues involved in the cavity should have an RMSF lower than 4.1 nm2. The latter criterion is motivated by the fact that we require a binding site to be sticky, since the aim of this study is to bind sticky residues with small molecules which in turn aims to inhibit phase separation caused by interactions between sticky residues as noted by^20^. This procedure results in the identification of 41 binding sites throughout the ensemble of AR-V7.

### 3.3 Molecular Docking

For the screening of the compounds we randomly sampled a list of 6000 compounds from Chembl database^22^. We created pdbqt files from the smile presentation of the molecule using Ligpreper suite with and we started the docking against the selected binding sites. For each of the 41 binding sites of the AR-V7 structural ensemble we perform molecular docking simulation of each compound using Autodock Vina, an established open source software that can screen molecules with a relatively good reported accuracy^30^. After the docking against the 41 identified binding sites, using a 16 cpu machine, we gather the molecules into a single dataset containing the smiles and the predicted energy of each molecule for each binding site of the protein. The overall dataset consists of 6000 molecules screened against these 41 sites summing to the total 246000 available energies.

### 3.4 Machine learning

In order to efficiently screen a vast space of compounds, using the molecular docking data of the previous step, we train a multi label classification model that predicts the activity of a compound against any of the 41 binding sites. The dataset is created as follows. First, we identify the binding energy threshold of the top 5% binding compounds for each of the 41 binding sites. Then, –for each binding site– a compound is rendered as active “1” if the docking energy is lower than the energy threshold of that binding site and “0” otherwise. Overall, there are 41 classification endpoints for each compound. We attained to train an efficient model for the 5% for the top energies by oversampling the “active” molecules since the dataset is heavily imbalanced towards the non active molecules. The model trained on the top 5% of the energies is selected to screen molecular libraries. For the multi label classification model (QSAR), the best model was trained using Topological Fingerprints as embedding generated using RDKit and a plain feed forward neural network. The topological fingerprints were of the size of 1412. The QSAR classification model is a feed forward neural network with 4 layers of size 1256. We used the rectified linear unit (ReLU) activation function, added a dropout on the first and the final layer, to avoid overfitting. In the output layer we used a sigmoid function. For the loss calculation we used a binary cross entropy loss (eq. 1) which is the loss function typically used in classification and multi label classification models. The validation of the model takes place on the 10% of the dataset. We splitted the molecules in the train dataset consisting of 90% of the molecules and the test with the rest 10% of the molecules. We created the descriptors directly from the smiles of the sampled molecules. For the inference of the model we stored the model with the best overall mean Matthews correlation coefficient (eq. 2). The MCC value was calculated per class. The methodology consists of the following steps:

1. Retrieving the conformational ensemble of the biomolecular target of interest. In this study we use AlphaFold-Metainference ^12^ due to speed and accuracy efficiency reasons.
2. Pocket detection and filtering down the selected binding sites. This part of the methodology can vary between proteins, due to the biological function of the protein that needs to be altered. In the case of AR-V7, in an effort to inhibit aberrant liquid-liquid phase separation, which in turn leads to aberrant transcription activation and tumor growth, we aim at detecting binding sites around sticky residues, since such residues are responsible for formation liquid-liquid phase separation.
3. Small molecule list selection and molecular docking of this list against the detected cavities. Here we used randomly sampled molecules from the Chembl database. Other criteria such as Lipinski rule of 5, could be used to subsample efficiently the chemical space.
4. Using data created in 3., create a multilabel classification QSAR model and accelerate the screening procedure by inferring a vaster chemical space by using the QSAR model generated. In the specific case we accepted as good predictions the molecules that predicted as active “1” more than 70%, hence they interact with more than 70% of the docking sites. The threshold can vary from protein to protein and screening results. The molecules passing the threshold are chosen for re-docking.

### 3.5 Evaluation metrics

We evaluated the models for both multiclass classification and per docking site. For the multiclass we used the evaluation metrics of zero one loss and the hamming loss. For the evaluation of the predictions for each docking site we used the Matthews Correlation Coefficient (MCC).

Binary cross entropy, also known as Binary Log Loss or Binary Cross-Entropy loss, is a commonly used loss function in machine learning. It is mainly used in binary classification problems and is designed to measure the dissimilarity between predicted probability distribution and the true labels of a dataset.

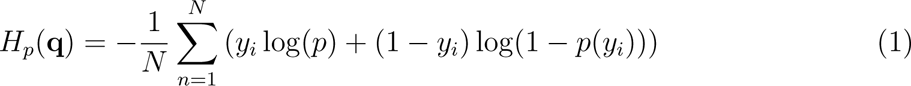

The MCC can be quantified as in (eq. 3), where tp quantifies the true positives per binding site tn are the true negatives per binding sites fp is the false positives per docking site fn are the false negatives and is essentially a measure of the quality of binary and multiclass classifications ^31^.

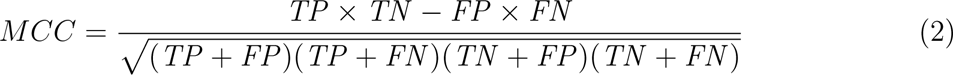

The Zero One Loss can be quantified as in (eq. 1), where y are the truth labels of the active / non active and *y^i^* are the predicted classes and *n_samples_* is the number of the samples.

In multilabel classification, the zero one loss corresponds to the subset zero-one loss: for each sample, the entire set of labels must be correctly predicted, otherwise the loss for that sample is equal to one^32^.

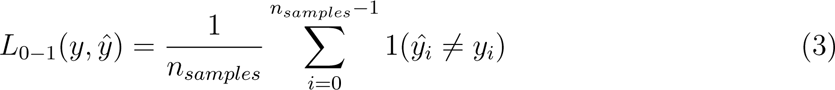

The Hamming Loss can be quantified as in (eq. 2), where y are the truth labels of the active / non active and *y*^ are the predicted classes and *n_samples_* is the number of the samples. The Hamming loss is the fraction of labels that are incorrectly predicted^32^.

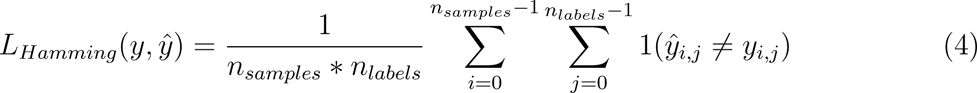

